# Antagonism of Saxitoxin and Tetrodotoxin Block by Internal Monovalent Cations in Squid Axon

**DOI:** 10.1101/2021.12.21.473550

**Authors:** Gerry S. Oxford, Paul Forscher, P. Kay Wagoner, David J. Adams

## Abstract

The block of voltage-dependent sodium channels by saxitoxin (STX) and tetrodotoxin (TTX) was investigated in voltage-clamped squid giant axons internally perfused with a variety of permeant monovalent cations. Substitution of internal Na^+^ by either NH_4_^+^ or N_2_H_5_^+^ resulted in a reduction of outward current through sodium channels under control conditions. In contrast, anomalous increases in both inward and outward currents were seen for the same ions if some of the channels were blocked by STX or TTX, suggesting a relief of block by these internal cations. External NH_4_^+^ was without effect on the apparent magnitude of toxin block. Likewise, internal inorganic monovalent cations were without effect, suggesting that proton donation by NH_4_^+^ might be involved in reducing toxin block. Consistent with this hypothesis, decreases in internal pH mimicked internal perfusion with NH_4_^+^ in reducing toxin block. The interaction between internally applied protons and externally applied toxin molecules appears to be competitive, as transient increases in sodium channel current were observed during step increases in intracellular pH in the presence of a fixed STX concentration. In addition to these effects on toxin block, low internal pH produced a voltage-dependent block of sodium channels and enhanced steady-state inactivation. Elevation of external buffer capacity only marginally diminished the modulation of STX block by internal NH_4_^+^, suggesting that alkalinization of the periaxonal space and a resultant decrease in the cationic STX concentration during NH_4_^+^ perfusion may play only a minor role in the effect. These observations indicate that internal monovalent cations can exert trans-channel influences on external toxin binding sites on sodium channels.

## INTRODUCTION

The movement of sodium ions through voltage-dependent sodium (Nav) channels during excitation of nerve and muscle is selectively and reversibly inhibited by tetrodotoxin (TTX) and saxitoxin (STX). These compounds have proven crucial to our understanding of the behaviour of both Na and K channels, to determinations of the surface density of Nav channels, and are employed in the biochemical isolation, characterization, and functional reconstitution of Nav channels (for reviews see Narahashi, 1974; Catterall, 1992, 1995). Equilibrium and kinetic studies of toxin-receptor for interaction indicate that one toxin molecule blocks one Nav channel (Hille, 1968; Cuervo and Adelman, 1970; Keynes et al., 1971; Colquhoun and Ritchie, 1972). Although it has been proposed that positively charged guanidinium groups on the toxin molecules enter and bind to a site within the channel lumen to block ion flow (Kao and Nishiyama, 1965; Hille, 1975), recent evidence suggests that these toxins bind to a receptor on the external surface of the channel close to, but not in, the permeation pathway (Kao and Walker, 1982; Strichartz, 1984).

Toxin block has been reported to be independent of the species of external cation (Moore et al., 1967) or the direction of current flow (Rojas and Atwater, 1967), yet the binding of radiolabelled TTX and STX is competed by monovalent cations which permeate open channels (Henderson et al., 1974; Barchi and Weigele, 1979; Strichartz et al., 1986). In addition, TTX and STX binding is influenced by extracellular pH (Colquhoun et al., 1972; Henderson and Wang, 1972), external divalent cations (Hille et al., 1975; Strichartz et al., 1986), and solvent properties (Hahin and Strichartz, 1981). Using electrophysiological methods, Ulbricht and Wagner (1975) confirmed that protons compete with TTX for the same site and that occupation of the site either by protons or by toxin would block the Nav channel. In contrast, the influence of divalent cations on toxin effectiveness has been attributed to surface potential induced changes in effective toxin concentration rather than ion-toxin competition (Hille et al., 1975; Grissmer, 1984; but see Strichartz et al., 1986).

The present investigation stems from an observation made during previous experiments on K channel properties (Oxford and Adams, 1981) that very small outward transient currents would consistently appear in voltage-clamped squid axons bathed in TTX when the internal K^+^ ions were substituted by NH_4_^+^ ions as charge carriers. The kinetics of these transient currents were reminiscent of Nav channels and they could be reduced by conditioning depolarizations. We thus sought to define more clearly the interactions between permeating cations and the toxin binding site and to investigate the possibility that outward ion movement could modulate the binding of toxin applied to the external surface. A preliminary report of these results has been published (Oxford, Forscher and Adams, 1984).

## MATERIALS AND METHODS

Experiments were performed on internally perfused giant axons isolated from *Loligo pealei* at the Marine Biological Laboratory, Woods Hole, MA. The axons were voltage-clamped using conventional axial electrode techniques. Details of the electronics, perfusion, and associated procedures for voltage clamping have been published (Oxford et al., 1978; Oxford, 1981). Ionic currents were low pass filtered (Bessel) at 30 kHz and recorded via a 12-bit A/D convertor interfaced to a PDP 11/23 computer for data storage and analysis. Capacitative and linear leakage currents were digitally subtracted from the records using a “-P/4” averaging procedure. The holding potential was maintained at −80 mV in all experiments and compensation for series resistance was employed as detailed previously (Oxford, 1981).

The compositions of experimental solutions are given by the following notation: EXTERNAL // INTERNAL, and are detailed in Table I. Variations of the internal Na^+^ concentration were achieved by equimolar substitution of Cs^+^ (see Oxford and Yeh, 1985). Ionic current contributions through voltage-dependent K channels were reduced by internal 1 mM 3,4-diaminopyridine (DAP) (Kirsch and Narahashi, 1978). All experiments were done at a temperature of 10 ± 0.5°C controlled by a Peltier cooling device and feedback circuitry.

**TABLE 1.** Composition of External and Internal Solutions

| **External Soln.** | **Na** | **TMA** | **Tris** | **NH<sub>4</sub>** | **Ca<sup>2+a</sup>** | **C1** | **HEPES<sup>b</sup>** |
| --- | --- | --- | --- | --- | --- | --- | --- |
| 450 Na// | 450 |  |  |  | 50 | 545 | 5 |
| 450 TMA,5 HEPES// |  | 450 |  |  | 50 | 545 | 5 |
| 450 TMA,25 HEPES// |  | 410 |  |  | 50 | 510 | 25 |
| 450 Tris// |  |  | 450 |  | 50 | 545 | 5 |
| 450 NH<sub>4</sub>// |  |  |  | 450 | 50 | 545 | 5 |

| **Internal Soln.** | **Na** | **Cs** | **F** | **Glucose** | **Sucrose** | **MOPS<sup>d</sup>** |
| --- | --- | --- | --- | --- | --- | --- |
| //300 Na<sup>c</sup> | 300 |  | 50 | 250 | 345 | 10 |
| //100 Na,200 Cs | 100 | 200 | 50 | 250 | 345 | 10 |

TMA, tetramethylammonium; Tris, tris(hydroxymethyl)aminomethane; HEPES, 4-(2-hydroxyethyl)-1-piperazineethanesulfonic acid; MOPS, 3-(*N*-morpholino)propanesulfonic acid; MES, 2-(*N*-morpholino)ethanesulfonic acid.
<sup>a</sup> In one series of experiments 10 mM CaCl<sub>2</sub> and 40 mM MgCl<sub>2</sub> were substituted. No significant differences were observed by this substitution.
<sup>b</sup> External solutions were buffered to pH 7.8.
<sup>c</sup> Equimolar substitutions of monovalent cations (Li<sup>+</sup>, Cs<sup>+</sup>, NH<sub>4</sub><sup>+</sup>, N<sub>2</sub>H<sub>5</sub><sup>+</sup>, and K<sup>+</sup>) were made for Na<sup>+</sup>.
<sup>d</sup> pH was adjusted to 7.3 for most experiments. 10 mM MES (pK<sub>a</sub> 6.2) was substituted for MOPS (pK<sub>a</sub> 7.2) in experiments where the pH was reduced to 5.6.

Internal and external perfusion rates were 50-100 μL,/min. and 0.3 ml/min., respectively. Given the volumes of the chamber and axons, this corresponds to solution exchange times of 0.4 and 20 sec for the intracellular and extracellular compartments, respectively.

The concentrations of saxitoxin (STX) and tetrodotoxin (TTX) used in these experiments are overestimates as no correction was made for the degradation of toxin stocks. Chemical reagents used were of analytical grade and 3,4-diaminopyridine was obtained from Aldrich Chemical Co. (Milwaukee, WI).

## RESULTS

### Outward ammonium currents are anomalously large at low STX doses

Under normal intracellular ionic conditions, potassium ions carry most of the outward current through Nav channels, thus we first sought to examine STX block of outward K^+^ currents. In Figure 1 outward K^+^ current families are shown for an axon bathed in 450 mM TMA, 5 mM HEPES// and internally perfused with 300 mM K^+^ and 1 mM DAP under control conditions (Fig. 1A) and after the addition of 10 nM STX to the external solution (Fig. 1B). Transient K^+^ currents through Nav channels were reduced 74% by STX. The axon was then returned to STX-free 450 TMA, 5 HEPES// and internally perfused with 300 mM NH_4_^+^. Following recovery of the transient outward currents, now carried by NH_4_^+^ (Fig. 1C), 10 nM STX was again introduced in the external medium, but in this case, it reduced the transient current by only 12% (Fig. 1D). This suggests that NH_4_^+^ currents through Nav channels interfere with the action of STX.

**Figure 1.**
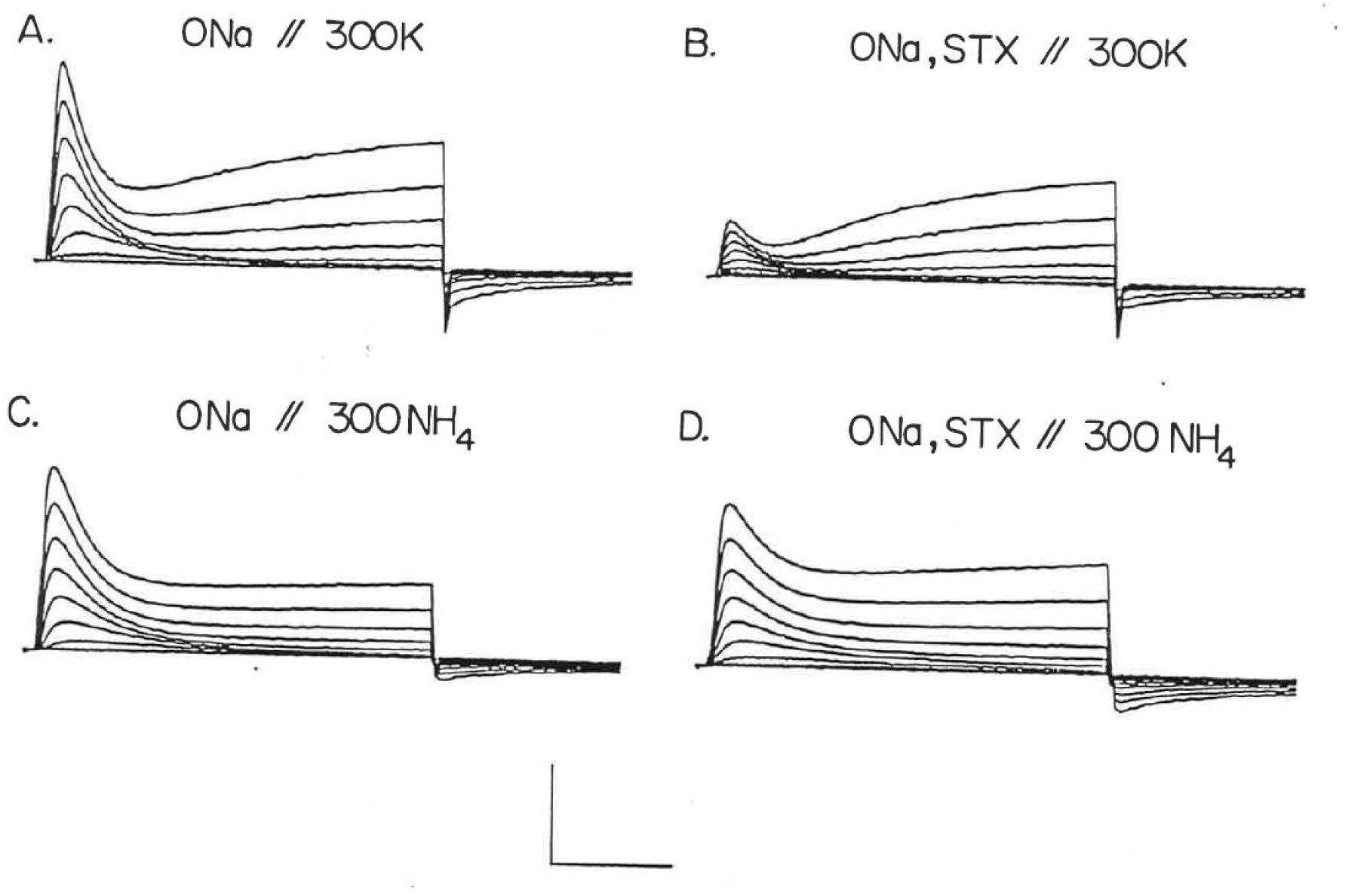
Internal ammonium perfusion reduces STX block of K^+^ currents through sodium channels. **(A)** Potassium currents through sodium channels (and potassium channels) in the absence of STX. **(B)** Currents from the same axon as (A) following the addition of 10 nM STX to the bathing medium. Note the selective reduction of sodium channel current. **(C)** Outward ammonium currents following washout of STX and substitution of internal K^+^ by NH_4_^+^. **( D)** Reintroduction of 10 nM STX to the external bath has very little effect on the amplitude of NH_4_^+^ currents through the sodium channels. Scale markers represent 0.36 mA/cm^2^ and 2 ms (A and B), or 0.72 mA/cm^2^ and 2 ms (C and D). Temperature = 10.2°C.

We sought to compare the efficacy of STX block of Nav channels for various permeant cations. In order to assess toxin block for ionic currents of comparable magnitude, we first established the current-carrying capacity of different ions through Nav channels. The magnitudes of outward ionic currents at +100 mV are illustrated in Figure 2 (solid bars) for various internal monovalent cations and concentrations normalized to values in 300 mM Na^+^. The magnitude of outward current increases non-linearly for increases in Na^+^ concentration (Hille, 1975). Since comparable currents were obtained for [Na^+^] near 100 mM and [NH_4_^+^] at 300 mM, we compared STX block in these two cases. In Figure 3, current families from one axon bathed in 450 TMA, 5 HEPES and internally perfused alternately with //100 Na^+^ and //300 NH_4_^+^ are compared under control conditions (Fig. 3A,B) and following addition of 20 nM STX (Fig. 3C,D). It is clear that 20 nM STX produces much more block of outward Na^+^ currents (Fig. 3C) than of outward NH_4_^+^ currents (Fig. 3D).

**Figure 2.**
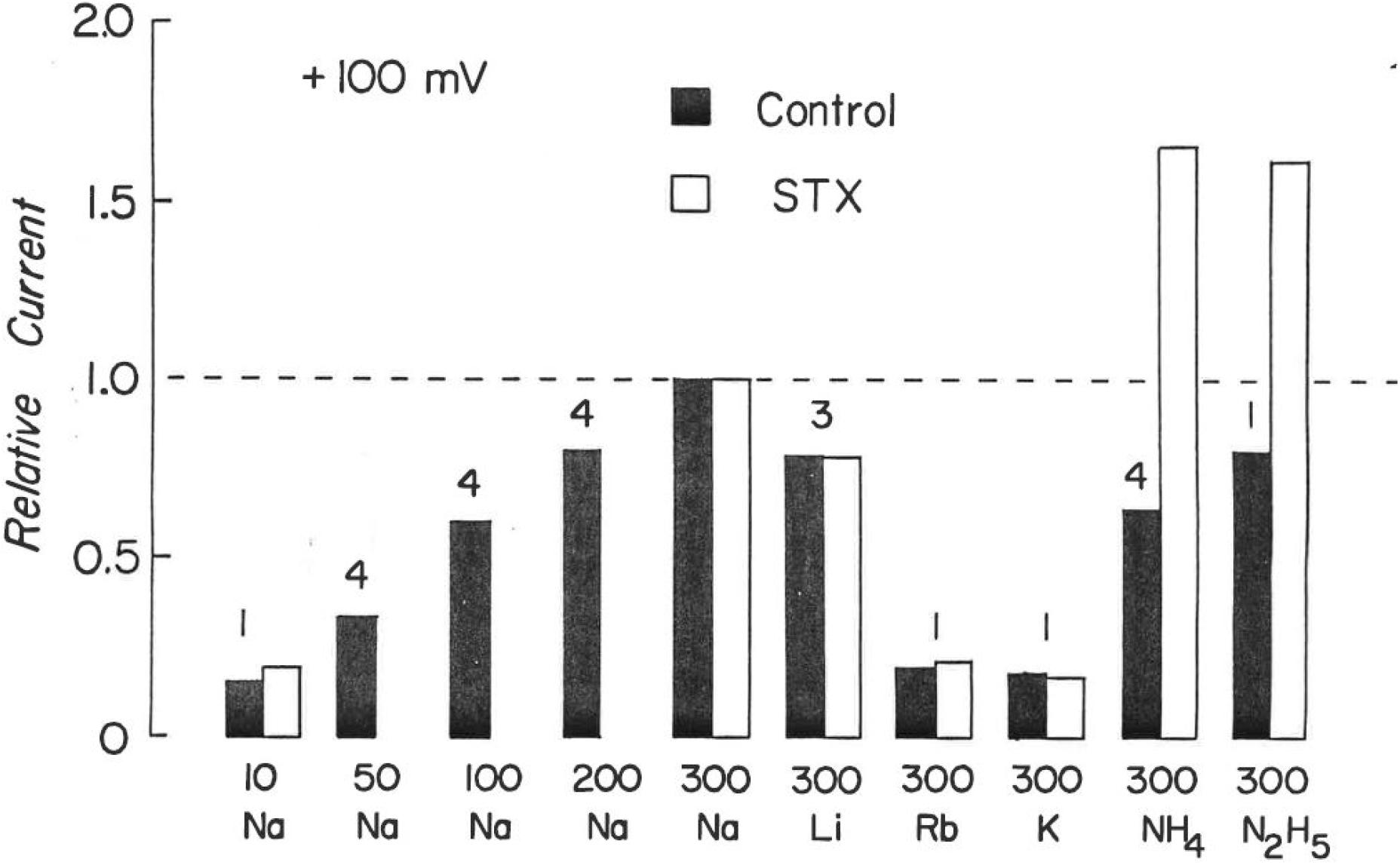
Magnitude of outward current through sodium channels for various internal monovalent cations in the presence and absence of 10-20 nM STX. Solid bars represent mean peak outward currents at +100 mV for the indicated internal cations and concentrations (mM) normalized to the values obtained in the same axons at 300 mM Na^+^. Open bars represent similar data obtained in the presence of either 10 or 20 nM STX and again normalized to the current obtained in 300 mM Na^+^. Numbers over each set of bars correspond to the number of experiments for both toxin and control conditions.

**Figure 3.**
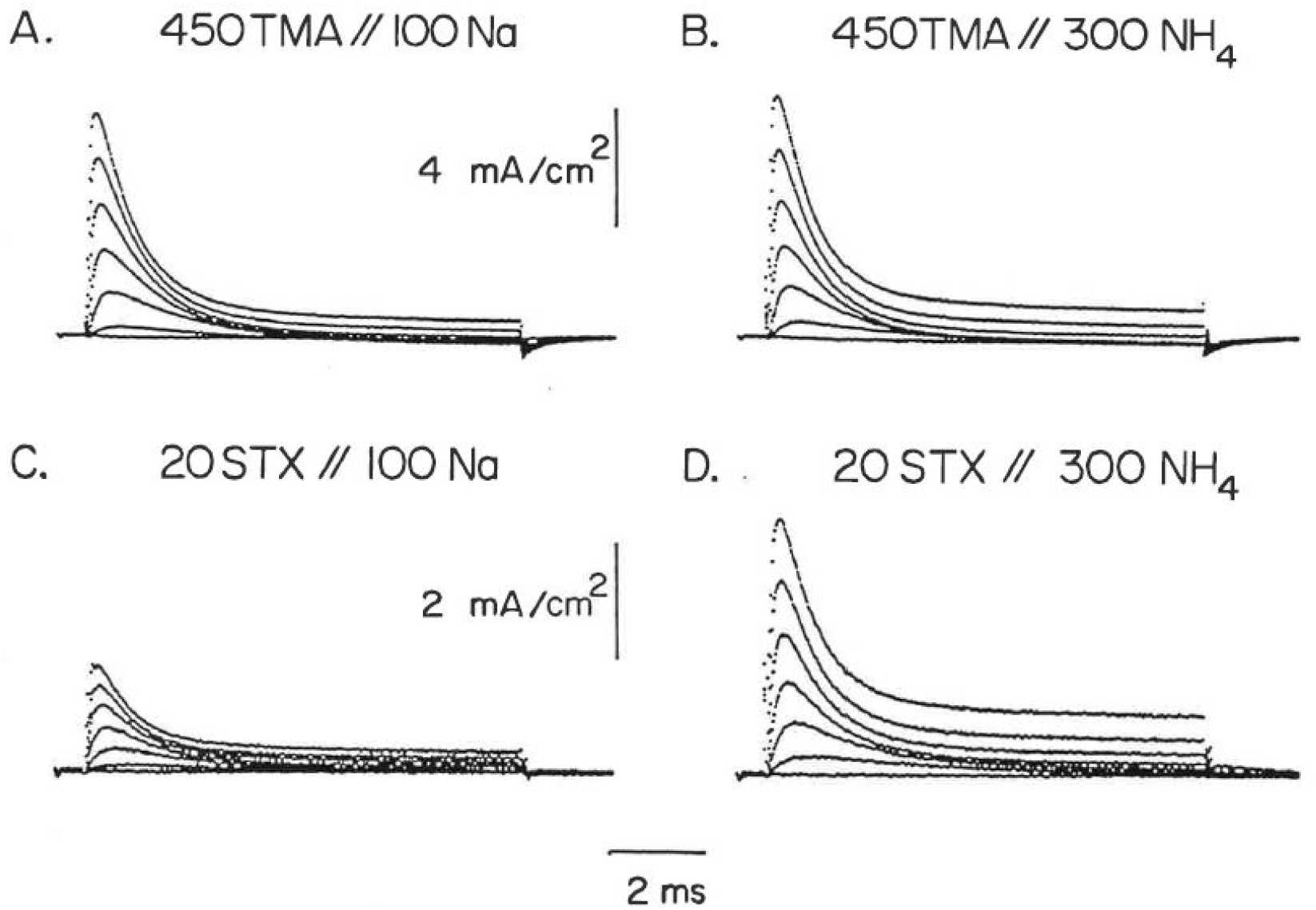
Comparison of STX block in Na^+^-perfused and NH_4_^+^-perfused axon exhibiting comparable current magnitudes. **(A)** Outward sodium currents in an axon perfused with //l00 Na. **(B)** Outward ammonium currents from the same axon following perfusion with //300 NH_4_. **( C)** Outward currents following return to //l00 Na and addition of 20 nM STX to the external 450 TMA solution. **(D)** Currents in the presence of STX during perfusion with //300 NH_4_. Note the change in vertical scales between A,B and C,D.

Evaluation of outward ionic currents for the various monovalent cations in the presence of 20 nM STX (Fig. 2, open bars) reveals that of the ions tested, only ammonium and hydrazinium ions produced anomalously large currents when compared to currents in the absence of toxin. Thus, these ions apparently weaken the block of sodium channels by STX at this concentration. Further experiments exploring this phenomenon were addressed with NH_4_^+^ ions as the axons were more stable when perfused with ammonium solutions than with hydrazinium solutions.

### External NH_4_^+^ does not reduce STX block of Nav channels

Ammonium ions can carry current through Nav channels in either direction (Binstock and Lecar, 1969). We thus examined whether the interaction of NH_4_^+^ ions with Nav channels during inward currents also resulted in an apparent reduction of toxin block. The degree of toxin block was represented as the ratio of ionic currents in STX vs. control and is plotted as a function of membrane potential in Figure 4A for two axons bathed in 450 Na^+^// (closed symbols) and then 450 NH_4_^+^// (open symbols). Substitution of external NH_4_^+^ for Na^+^ did not alter the degree of block by STX for inward currents at potentials more negative than the reversal potential (measured as +33 mV in both cells, see legend). In addition, outward currents carried by Na^+^ ions for both cases were also equally sensitive to STX. Thus unlike the case for internal NH_4_^+^ (Fig. 4B, see below), neither the presence of extracellular NH_4_^+^, nor its inward movement through the channel, altered the action of STX on Nav channel currents in either direction.

**Figure 4.**
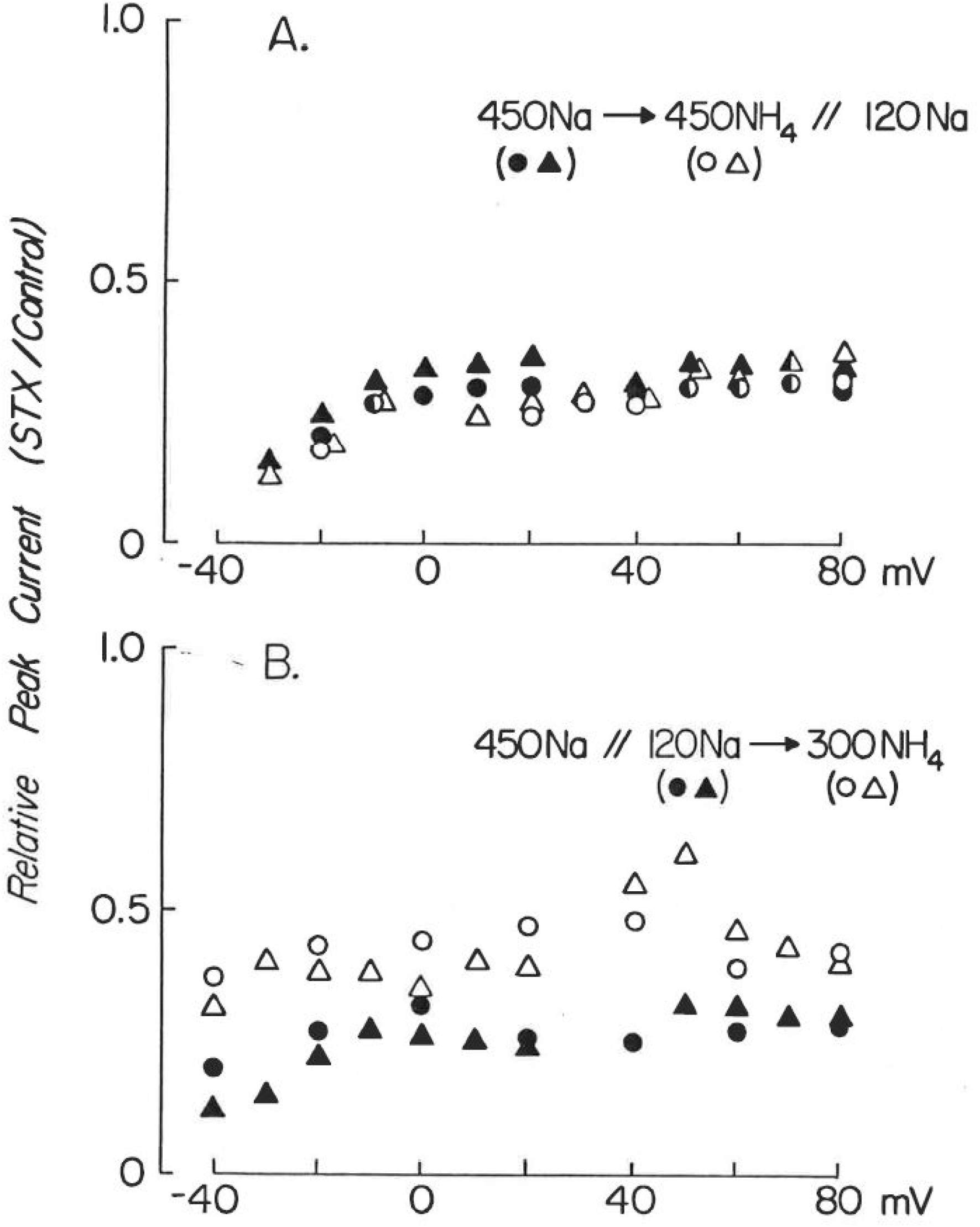
Relative peak sodium channel currents following exposure to 20 nM STX as a function of membrane potential. Data points represent the ratio of peak current in STX to that in the absence of toxin. **( A)** External ammonium ions (open symbols) do not affect STX block compared to that seen with 450 mM Na (closed symbols). Circles and triangles represent 2 separate axons. **( B)** Internal ammonium perfusion reduces STX block of inward sodium currents. Closed symbols are from 2 axons (different from A) perfused with //120 Na and then perfused with //300 NH_4_ (open symbols). Reversal potential observed for both conditions was +33 mV.

### Internal NH_4_^+^ inhibits STX block of inward Na^+^ currents

To test the effect of internal NH_4_^+^ on inward Na^+^ currents in the presence and absence of 20 nM STX, 300 NH_4_^+^ was exchanged for 120 Na^+^ in the internal perfusate (Fig. 5). Although inward currents in //120 Na^+^ and //300 NH_4_^+^ measured in the absence of toxin were identical as were the reversal potentials, inward Na^+^ currents during STX exposure increase when NH_4_^+^ (Fig. 5B) replaces Na^+^ (Fig. 5A) internally. These observations on 2 axons are summarized in Figure 4B for a range of membrane potentials during internal perfusion with //120 Na^+^ (closed symbols) and then //300 NH_4_^+^ (open symbols). It thus appears that inhibition of toxin block by internal NH_4_^+^ is not limited to ion movement in the outward direction, and that occupancy of an open Nav channel by NH_4_^+^ may not be required for the effect on STX block.

**Figure 5.**
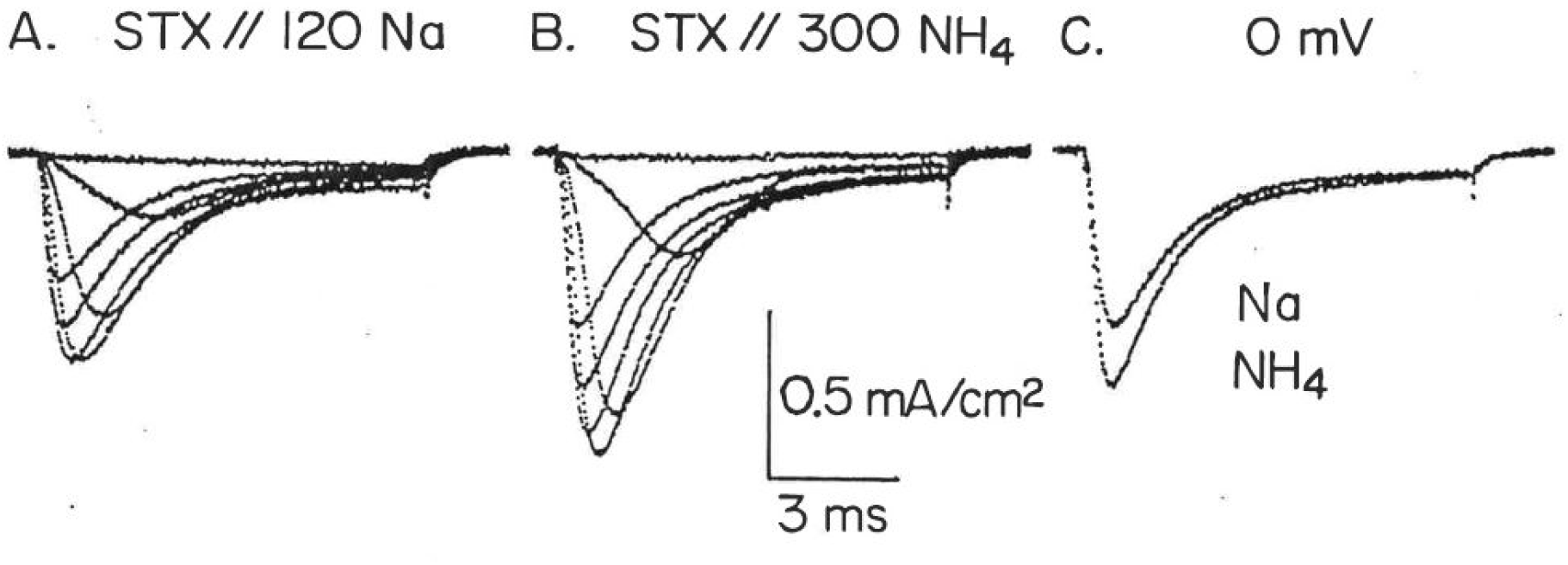
Internal ammonium increases inward sodium currents in the presence of STX. Ionic current responses to membrane potential steps to −50, −40, −30, −20, −10, 0, and +10 mV in an axon exposed to 20 nM STX in 450 Na and internally perfused first with //120 Na **(A)** and then //300 NH_4_ **(B)** A comparison of the records at 0 mV is shown in C, where the larger current was obtained in //300 NH_4_.

### Ammonium ions inhibit TTX block of Nav channels

While STX and TTX exhibit identical actions on Nav channels, they are chemically different molecules in terms of structure, valence, and pKa. TTX is monovalent at physiological pH (pK_a_ = 8.8), whereas STX is divalent (pK_a_s = 8.25 and 11.6). Such differences might be reflected in different sensitivities of their block to inhibition by ammonium ions. To examine this question, we determined the degree of block by TTX in axons perfused with 100 Na^+^ as compared to 300 NH_4_^+^. Internal NH_4_^+^ ions reduced TTX block of Nav channel currents as they did for STX block. An example of this effect is illustrated in Figure 6, where comparisons of outward Na^+^ and NH_4_^+^ currents at +100 mV are made in the absence (Fig. 6A) and presence (Fig. 6B) of 10 nM TTX. While the peak current amplitudes are identical without toxin, in the presence of toxin NH_4_^+^ currents are approximately twice as large as Na^+^ currents.

**Figure 6.**
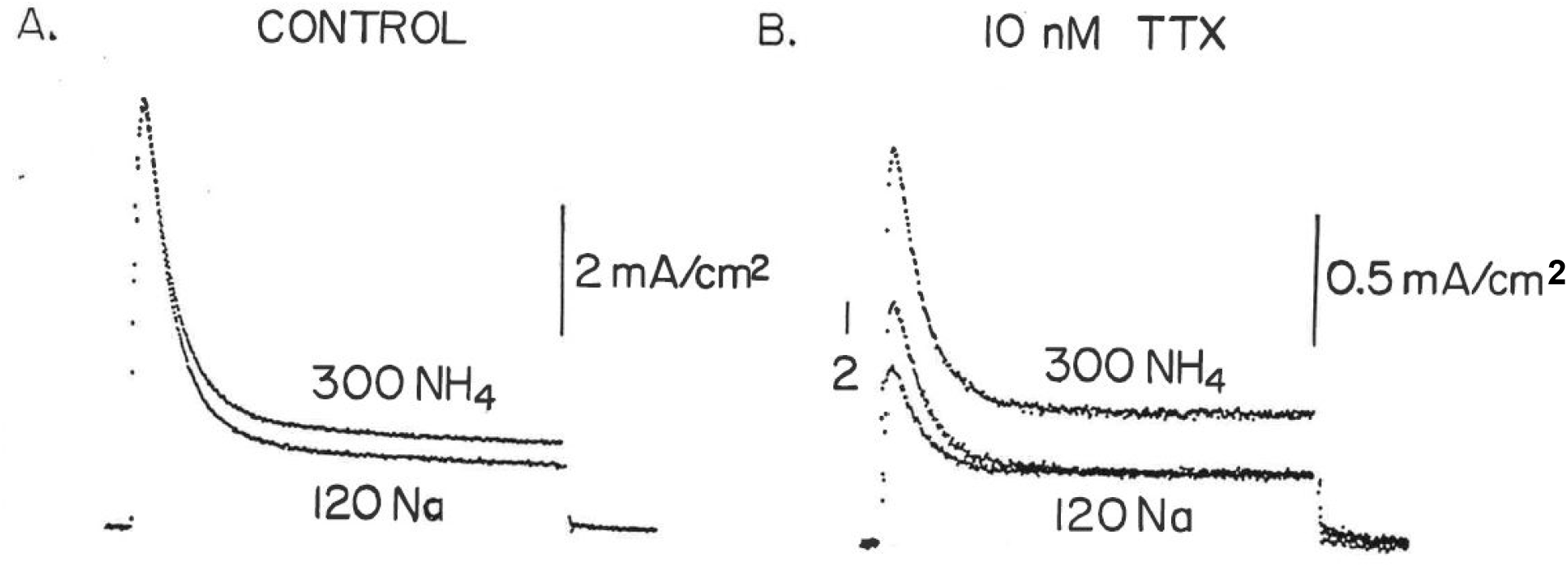
Ammonium perfusion reduces TIX block of sodium channels. **(A)** Outward currents at +100 mV for an axon perfused consecutively with //120 Na and //300 NH_4_. External solution was 450 TMA. Note the identity of peak current in the two solutions. **(B)** Outward currents under similar conditions following addition of 10 nM TTX to the bath. Records labelled 1 and 2 were obtained before and after perfusion with //300 NH_4_, respectively. Note the increase of outward current with ammonium perfusion. Increased outward steady-state current with //300 NH_4_ in both cases probably reflects NH_4_^+^ movement through K channels not blocked by DAP. Note the 4-fold difference in vertical scales between A and B.

### Internal protons inhibit Nav channel block by external STX and TTX

A common feature that distinguishes ammonium and hydrazinium ions from the other monovalent cations tested, is their ability to act as proton donors in aqueous solution. In view of the effectiveness of these cations in modulating STX block and of previous evidence that protons compete with STX and TTX for their receptor sites (Ulbricht and Wagner, 1975), we considered the possibility that proton donation by NH_4_^+^ and N_2_H_5_^+^ might be involved in inhibition of toxin action. To examine this point, we compared STX block of Nav channels as a function of intracellular pH. In the absence of toxin, we examined the effects of lowering intracellular pH on Nav channel currents. Our experiments lead to three basic observations illustrated by the records in Figure 7. First, in agreement with previous reports (Carbone *et al*., 1981), we observed a reduction in Na^+^ current as the pH was lowered from 7.3 to 5.6. This reduction was voltage-dependent as shown for low external pH (Woodhull, 1973; Begenisich and Danko, 1983) such that proton block was relieved for increasingly positive membrane potentials (Fig. 7A,C). Estimates of the effective electrical location (Woodhull, 1973) of the blocking site within the pore made from peak currents at potentials more positive than 0 mV yielded a value of 0.41 ± 0.07 (n = 3, S.D.) from the external membrane surface. A second observation was that the steady-state inactivation of Nav channels was more complete at pH 5.6 than at pH 7.3. This is in agreement with earlier reports (Brodwick and Eaton, 1978; Carbone et al., 1981). A third observation was that the block of Nav channels by internal protons could be partially relieved by increases in the buffer strength in the <u>extracellular</u> solution at a constant pH of 7.8. Raising the external concentration of HEPES from 5 to 25 mM, while maintaining the internal pH at 5.6, increased Na^+^ currents at all potentials (compare Fig. 7B to A and C). When the internal pH was returned to 7.3, changes in external buffer strength had no effect on Na^+^ current magnitude (Fig. 7D,E,F). This result is consistent with the suggestion that protons can pass through open sodium channels (Mozhayeva and Naumov, 1983) and that alterations in buffer strength may modulate a “proton gradient” within the Nav channel pore and effectively regulate the concentration of protons at blocking sites. These observations thus confirm that protons can block sodium channels at sites within the electric field when elevated on the intracellular side.

**Figure 7.**
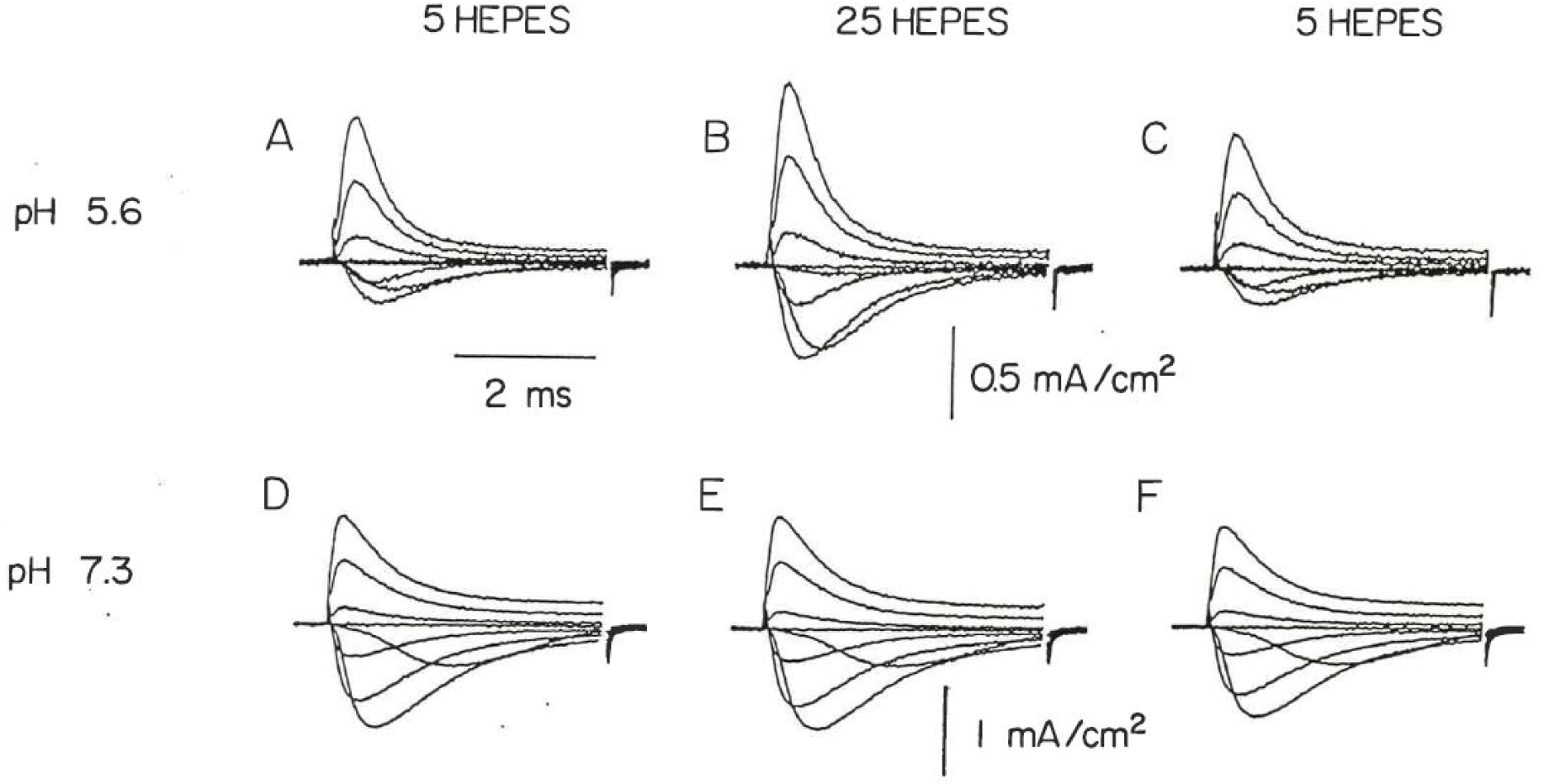
Increasing external buffer strength partially relieves block of sodium channels by internal protons. Families of sodium currents obtained consecutively in 450Na containing 5 mM HEPES, 25 mM HEPES, and return to 5 mM HEPES during perfusion with internal //l00Na at pH 5.6 (A,B,C) and then at pH 7.3 (D,E,F). Membrane potential steps to −60, −40, −20, 0, 20, 40, 60, and 80 mV from a holding potential of −80 mV.

We then compared the block of sodium channels by external 20 nM STX at internal pH values of 7.3 and 5.6. Equilibrium block as a function of membrane potential is illustrated in Figure 8 for 3 axons in the two pH conditions. Two observations can be made regarding these results. First, the effectiveness of STX on sodium channels is reduced by internal protons. The relative ionic current (STX/Control) at potentials above the reversal potential (+38 mV) is 0.29 ± 0.03 at pH 7.3, but increases to 0.46 ± 0.04 at pH 5.6. Second, the influence of internal protons on toxin block is progressively weaker for potentials more negative than the reversal potential. That is, the effect of protons on toxin block exhibits a voltage-dependence as does proton block alone.

**Figure 8.**
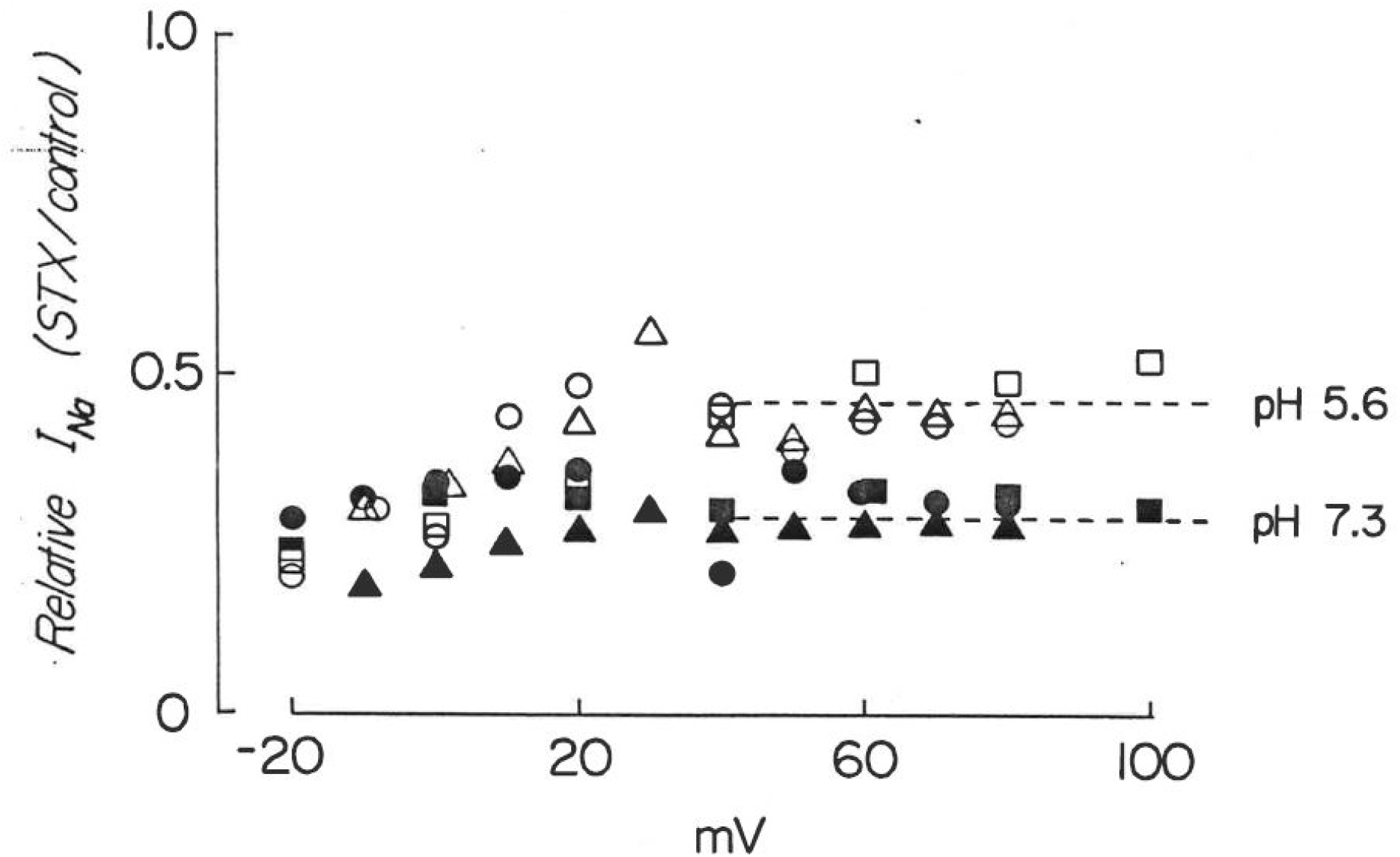
Low intracellular pH reduces block of sodium channels by external STX. Data points represent ratios of peak sodium currents obtained in the presence of 20 nM STX to those in the absence of toxin at several membrane potentials. Different shaped symbols apply to each of 3 axons, while corresponding closed and open symbols were obtained at internal pH values of 7.3 and 5.6, respectively. Dashed lines indicate the mean current ratios at pH 5.6 and 7.3 calculated from all data points above the reversal potential (+38 mV). Experimental solutions were 450 Na//100 Na for each axon.

### Internal protons and external toxin molecules interact in a competitive manner

Measurements of the kinetics of proton and toxin interaction with the sodium channel provide further support for an intimate relationship between their respective binding sites. In the absence of toxin, axons alternately perfused with solutions of pH 7.3 and 5.6 demonstrated monotonic increases and decreases in sodium current, respectively. The time constants for these changes in 2 axons were estimated to be 24 ± 8 seconds. As protonation/deprotonation reactions are rapid, this time probably reflects the solution switching time and associated dead space of the perfusion apparatus.

A different result was observed when the internal pH was switched during exposure to external toxin. Upon return from pH 5.6 to pH 7.3, the sodium current exhibited a <u>transient</u> increase. This phenomenon is illustrated in Figure 9 for an axon exposed to 20 nM STX. Current magnitudes at 0 mV were obtained at various time points before and after the switch of internal solutions from pH 5.6 to 7.3, and families of currents obtained at the times indicated by the letters. Sodium current increased rapidly as pH 7.3 solution entered the axon but then declined over several minutes to half of the maximum value. Examination of the current family records reveals two major additional points. First, as indicated previously following the switch to pH 7.3, sodium channel inactivation became less complete both at maximum current (**b**) and after several minutes of perfusion (**c**). Second, although the outward Na^+^ currents are of comparable magnitude in pH 5.6 (a) and following equilibration in pH 7.3 (**c**), the inward currents are smaller in pH 5.6. These observations suggest that the reduced currents in (**a**) (pH 5.6) reflect predominantly proton block which alters inactivation and is voltage-dependent, whereas the reduced currents in (**c**) reflect block by STX. The transient increase between the two measurements is expected if internal protons and external toxin compete for a common site (Ulbricht, 1981). As illustrated for 4 axons in Figure 10, such transients were seen in several experiments under similar conditions.

**Figure 9.**
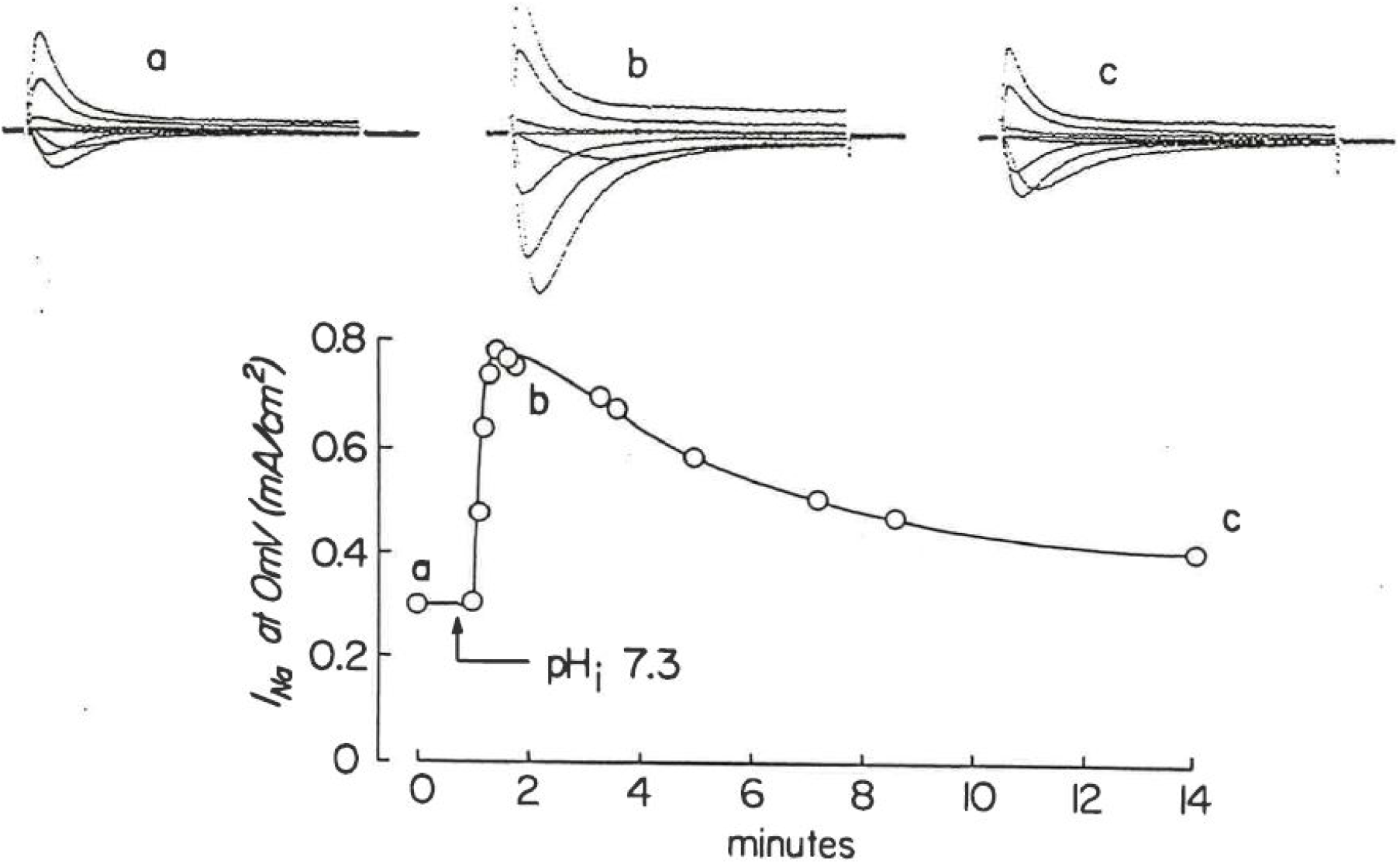
Peak sodium current measured at 0 mV at various times following a change in internal pH from 5.6 to 7.3 in the presence of external 20 nM STX. The data are plotted as a function of time (open circles) and reveal a transient recovery of current magnitude. Solid line fit by eye through the data points. Families of currents at several membrane potentials were obtained at the times indicated by the letters (a,b,c) and are depicted in the upper panel. Experimental solutions were 450 Na//100 Na.

**Figure 10.**
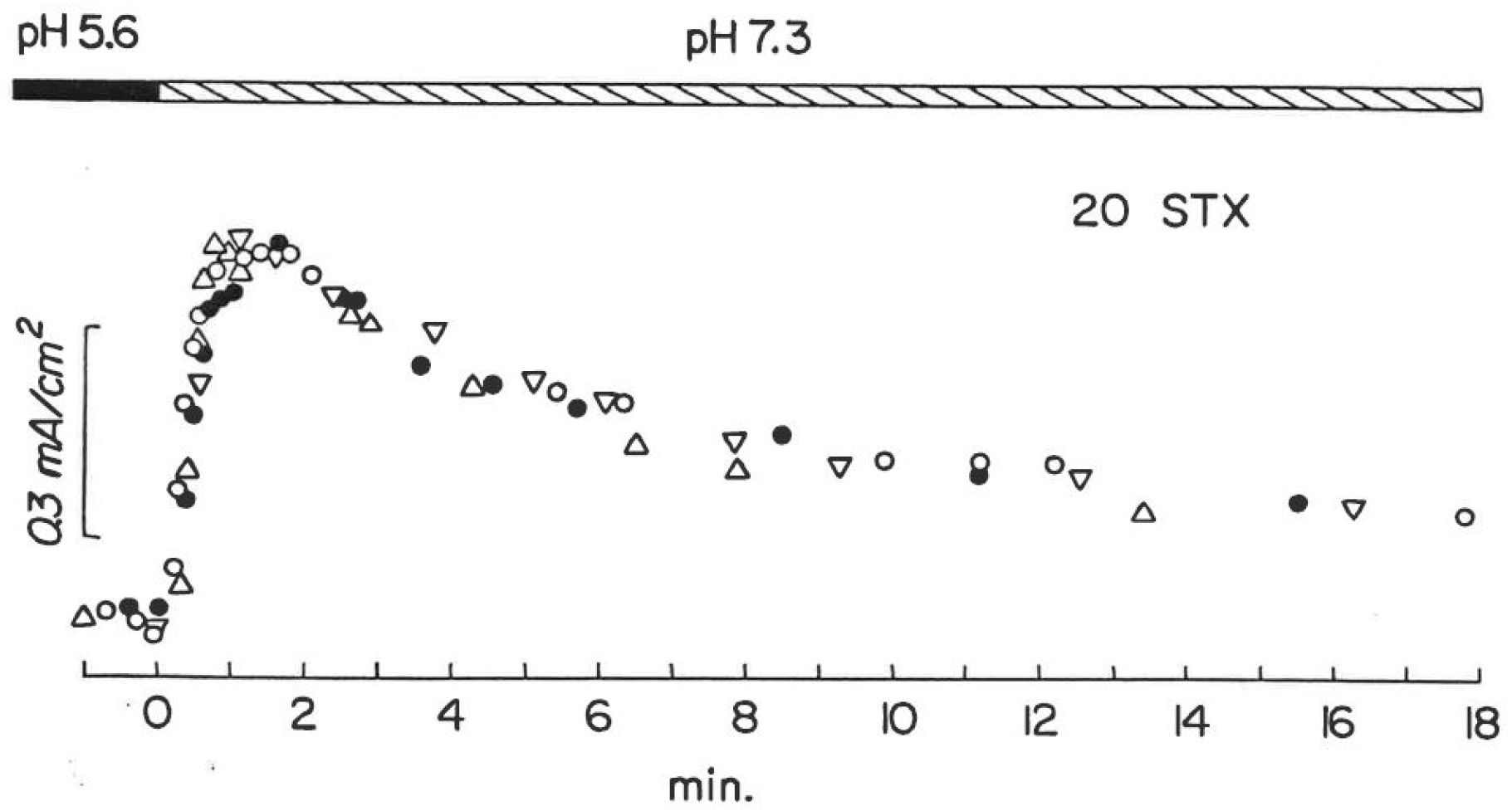
Sodium current transients recorded at 0 mV in 4 axons at various times during the transition between internal pH 5.6 and 7.3. Procedures and solutions identical to those of Figure 9. Data from each experiment has been shifted along the time axis to the estimated time of internal solution exchange at the recording region of each axon (0 min.).

The simplest interpretation of these transients is that protons and toxin molecules compete for a binding site, the occupation of which blocks current flow through the channel. At an internal pH of 5.6, the equilibrium for the competitive reaction is shifted towards protonation. When the internal pH is raised to 7.3, deprotonation rapidly occurs yielding unblocked channels (free toxin binding sites) and increases in current. Thereafter, a gradual reduction in current occurs due to the slower association of toxin with the channels. The magnitude of the transients reflects the ratio of the dissociation rate constants for protons and STX (Ulbricht, 1981).

### Is NH_4_^+^ relief of toxin block related to periaxonal alkalinization?

The similarities seen for sodium channel modulation by internal NH_4_^+^, N_2_H_5_^+^, and H^+^ ions supports the notion that proton donation by NH_4_^+^ and N_2_H_5_^+^ is involved in the antagonism of STX block. Such reactions by these cations could, however, lead to decreased STX block by a mechanism other than direct competition for a toxin binding site. External NH_4_Cl solutions have been extensively employed as a non-invasive means to elevate intracellular pH (Roos and Boron, 1981). In such cases, external NH_4_^+^ ions give up protons to the solution, the resultant NH_3_ molecules can diffuse into the cell, and there regain protons resulting in alkalinization of the intracellular compartment.

In the present experiments, it is conceivable that during axon perfusion with 300 mM NH_4_^+^, sufficient NH_3_ escapes to and accumulates in the periaxonal space beneath the Schwann cell layer resulting in an elevation of local extracellular pH. A sufficient alkalinization could result in deprotonation of the critical 7,8,9-guanidinium moiety (pKa 8.25, Kao et al., 1983) on some STX molecules, thus reducing the effective active toxin concentration. Clearly, a reduction in toxin concentration would yield less block during perfusion with internal NH_4_^+^.

As a test of this possibility, we determined the effect of NH_4_^+^ perfusion on STX block in axons bathed in solutions (pH 7.8) with elevated extracellular buffer concentrations. In our first experiments, we substituted 450 TMA with 450 Tris. In the 7 axons examined in 450 Tris, we never observed a relief of toxin block with internal NH_4_^+^ perfusion as seen routinely with 450 TMA. This result is consistent with the idea that Tris can buffer against periaxonal alkalinization which would otherwise reduce the active STX concentration. In contrast, in 3 axons, raising the concentration of HEPES buffer in 450 TMA from 5 mM to either 25 or 50 mM did not attenuate the relief of STX block by internal NH_4_^+^ perfusion. The relative effectiveness of 450 mM external Tris to 50 mM HEPES could simply reflect its higher concentration. However, the following experiment suggests an alternative explanation. In axons perfused with 120 mM Na^+^, exchanging external Tris for TMA results in a small suppression of Na^+^ conductance (Fig. 11A). In the presence of 20 nM STX, however, the currents in Tris are larger than those m TMA (Fig. 11B). This suggests that external Tris itself inhibits STX action (Fig. 11C) and may thus mask a relief of block by internal NH_4_^+^ perfusion.

**Figure 11.**
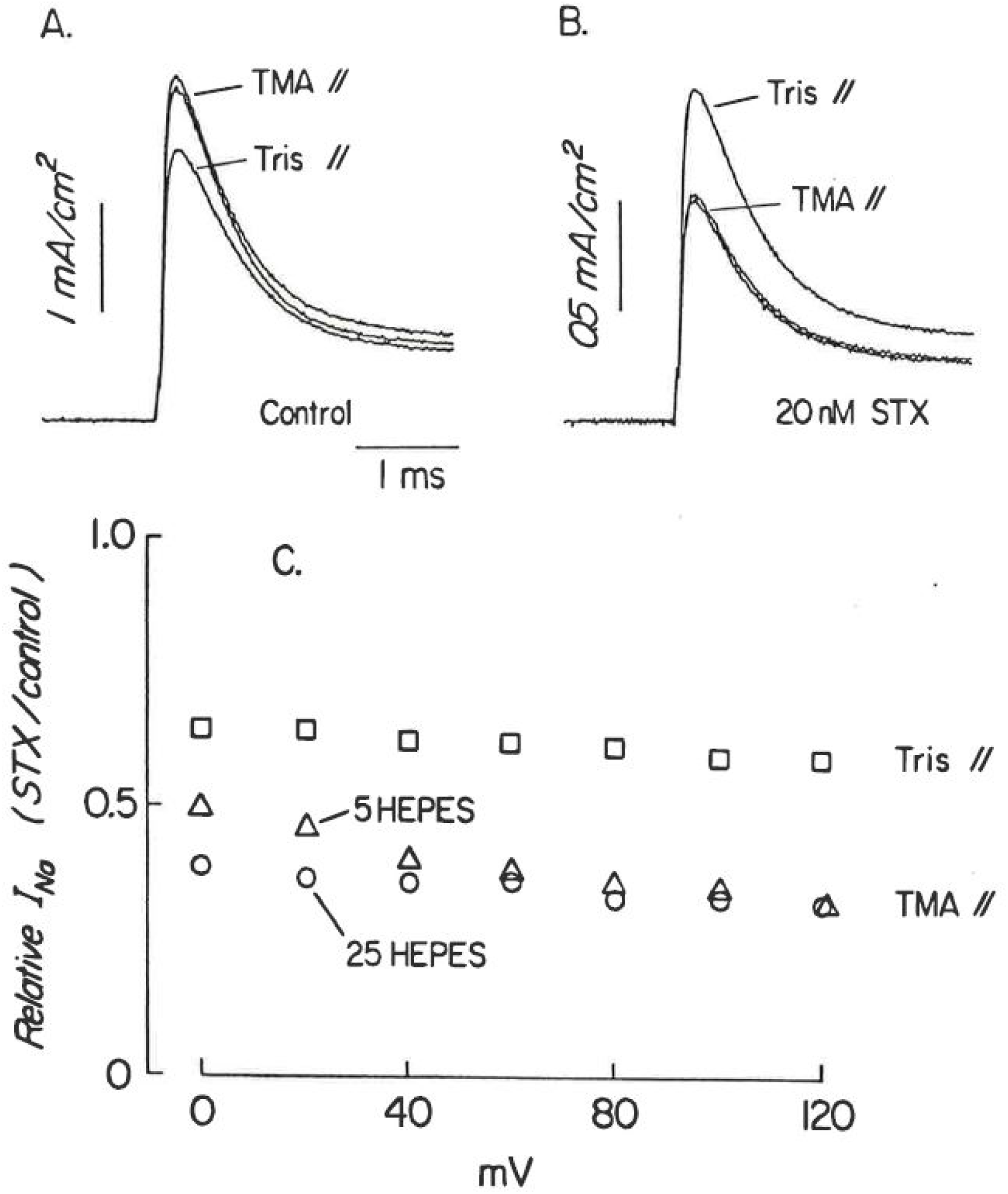
External Tris^+^ inhibits block of sodium channels by STX. Outward sodium currents were recorded consecutively at +100 mV in the following external solutions: 450 TMA, 5 HEPES//; 450 Tris//; and 450 TMA, 25 HEPES//. Measurements were made in the absence **(A)** and presence **(B)** of 20 nM STX. Note the increase in current with Tris^+^ in the presence of toxin. Ratios of current in toxin to control values are plotted in **(C)** for several membrane potentials in 450 TMA, 5 HEPES (triangles), 450 Tris (squares), and 450 TMA, 25 HEPES (circles). Internal solution was //120 Na.

As a second test for periaxonal alkalinization, we examined STX block at an external pH of 6.2. If alkalinization occurred during internal NH_4_^+^ perfusion, one would expect STX to be less susceptible to deprotonation at a pH further removed from its pKa of 8.25. In 2 axons examined, the relief of STX block by internal NH_4_^+^ was still present. However, the effect was significantly attenuated relative to that seen at an external pH of 7.8. Thus, the results from experiments to assess the influence of possible periaxonal alkalinization on the STX block neither conclusively demonstrate nor eliminate this alternative explanation.

## DISCUSSION

Block of sodium channels in squid giant axons by external STX and TTX is dependent on the species of internal monovalent cation. Specifically, internal protons and the organic cations, ammonium (NH_4_^+^) and hydrazinium (N_2_H_5_^+^) antagonize toxin block of ion movement through open Nav channels. In contrast, changes in the concentration of Na^+^ ions in the internal perfusate have no effect on toxin block. Consistent with this latter observation, it has been reported that toxin block of batrachotoxin-activated Nav channels in planar lipid bilayers is independent of internal Na^+^ concentration (Moczydlowski et al., 1984). Additionally, we have observed that the ratios of outward currents carried by various monovalent cations reflect an apparent conductance sequence of Na^+^ > Li^+^ = N_2_H_5_^+^ > NH_4_^+^ > Rb^+^ = K^+^. This conductance sequence is similar to the permeability sequence for these monovalent cations in Nav channels of squid axons (Chandler and Meves, 1965: Yamamoto et al., 1985). The conductance sequence for only the <u>metal</u> cations is independent of the presence of STX, whereas the sequence for all cations tested differs in the presence and absence of toxin (Fig. 2). It thus appears that the inhibition of toxin block by NH_4_^+^ or N_2_H_5_^+^ is not related to the relative permeability of these ions through the open unblocked Nav channel.

<u>External</u> monovalent metal cations have been shown to inhibit the binding of STX (Henderson et al., 1976; Weigele and Barchi, 1978) and TTX (Reed and Raftery, 1976: Frelin et al., 1981) in a manner which parallels their ability to impede ion flow through the Nav channel. However, recent STX binding studies in lobster nerves and membrane extracts of *Drosophila* brain (Strichartz et al., 1986) indicate that the dissociation constants for inhibition of Na^+^ permeability by <u>organic</u> cations and for organic cation competition of STX binding are different. Strichartz et al. (1986) conclude that the site at which these cations inhibit STX binding is distinct from the channel pore and selectivity filter.

Other evidence which argues against the site of toxin binding being the selectivity filter has been obtained from: (1) trimethyloxonium (TMO), a carboxyl group alkylating reagent modification of Nav channels to give TTX-resistant channels with normal ionic selectivity (Spalding, 1980), (2) structure-activity studies of TTX, STX and its analogues which suggest that the binding site is located at the orifice of the Nav channel (Kao and Walker, 1982), and (3) the difference between the dissociation constants for Na^+^ saturation of single channel conductance and for <u>external</u> Na^+^ competition of TTX block (Moczydlowski et al., 1984).

External divalent cations (e.g. Ca^2+^) also alter toxin binding and block of Nav channels, but in a manner different from that described here for internal monovalent cations. Divalent ion effects cannot be attributed to either direct competition or a simple action of the screening of fixed negative surface charges (see Grissmer, 1984; Strichartz et al., 1986). Modulation of toxin binding by divalent metal cations appears to be due to electrostatic interactions with negative surface charges in the vicinity of the toxin binding site (see Green and Andersen, 1986).

The antagonism of STX block by internal NH_4_^+^ does not appear to be a function of the direction of ion movement (Fig. 5; Rojas and Atwater, 1967). The reduction of STX block of inward Na^+^ current during internal NH_4_^+^ perfusion further suggests that occupancy of the Nav channel pore by NH_4_^+^ is unlikely to be required for the inhibition of STX block.

While external NH_4_^+^ does not impede toxin block, we did observe that replacing TMA^+^ with Tris^+^ inhibits STX action. Competition of STX binding by external organic cations may involve size-exclusion at the extracellular site such that higher dissociation constants are observed for larger organic cations. In this regard, we observed that the cation, Tris (M.W. = 121; 5.5 x 5.5 x 6.4 A) which is slightly larger than TMA (M.W. = 74; 5.5 x 5.5 x 5.5 Å), is more effective in competing with the STX molecule. The absence of effects of external NH_4_^+^ (M.W: = 18; 3.0 x 3.2 x 3.4 Å) on STX block (see Fig. 4A) is consistent with this notion.

### The involvement of protons in NH_4_^+^ antagonism of toxin block

The ability of ammonium ions to donate protons prompted an investigation of the effects of internal protons and their possible role in the NH_4_^+^ effect. Increasing the internal proton concentration (lowering pH) results in both block of the sodium channel and interference with the action of external STX and TTX. Both actions appear to reflect the ability of protons to pass through open sodium channels and interact with sites in the channel protein. In the absence of toxin, lowering the intracellular pH from 7.3 to 5.6 produced a voltage-dependent reduction in peak Na^+^ current, whereby the proton block was less at more positive membrane potentials (see also Wanke et al., 1980). The estimated fractional electrical distance for the blocking site is 60% of the channel length from the cytoplasmic side. This hypothetical locus compares roughly with the blocking site for external protons (Woodhull, 1973) estimated to be 25% of the way through the membrane field from the external surface.

Interactions between protons and other molecules associating with or entering sodium channels from the opposite side have been reported. Begenisich and Danko (1983) observed that raising the intracellular permeant cation concentration reduced the block of Nav channels by external H^+^, thus, demonstrating competition between the permeant cations and H^+^ for sites within the pore. We observed that the block of Nav channels by internal protons (pH 5.6) could be partially relieved by increasing the external buffer strength (HEPES concentration at a constant pH 7.8). This suggests that a proton gradient may exist through open Nav channels. Mozhayeva and Naumov (1983) directly measured proton currents through sodium channels in the node of Ranvier. Although the hydrogen to sodium permeability (P_H_/P_Na_) ratio they determined from reversal potential measurements equals 252, the channel conductance in a Na^+^-free acid solution is 50 times less than in control Na^+^ solutions. This suggests that protons can indeed ass through an open channel, but do so very slowly due to the binding to acidic group(s) in the channel.

Measurements of equilibrium block by STX at pH values of 7.3 and 5.6 (Fig. 8) indicate that the effectiveness of STX block is reduced by internal protons, such that there is a 63% relief of block (that is, increase of the relative Na^+^ current) at pH 5.6 compared to that observed at p 7.3. Relief of TTX block of peak Na^+^ permeability by external protons at the node of Ranvier has been previously reported and attributed to proton-toxin competition for the same site (Ulbricht and Wagner, 1975). If protons and toxin molecules interact with the same site on squid axon sodium channels, it may not be a site where protons alone block the channel. The slight voltage dependence of the proton-induced relief of toxin block is in the <u>opposite</u> direction to that of proton block alone suggesting that the locus at which protons block Nav channels and compete with toxin binding are different. The existence of multiple binding sites for protons within the Nav channel has been postulated from previous studies of H^+^ block of the Nav pore (Wanke et al., 1980; Begenisich and Danko, 1983).

The onset and recovery of Na^+^ current from block upon perfusing with internal solutions of pH 5.6 and 7.3, respectively, indicate that the time course of proton interaction with the Nav channel is a monotonic function. However, in the presence of toxin, a pH change from 5.6 to 7.3 results in a biphasic recovery of Na^+^ current characterized by a transient increase in both inward and outward Na^+^ current. The transient increase in Na^+^ current observed when the internal pH is raised from 5.6 to 7.3 may be attributed to the rapid dissociation of protons yielding unblocked channels which are blocked by STX with a slower association rate. As both protons and toxin molecules are positively charged, the apparent “competitive” interaction revealed by the transient increases in Na^+^ current may reflect mutual Coulombic exclusion rather than mutual affinity for the same site. Given that the toxin concentration at the external surface of the axon should have reached equilibrium during this experiment, the kinetics of the current decay after this transient increase should reflect the toxin-channel interaction rather than loading of the periaxonal space. The time constant to reach a new equilibrium at pH 7.3 (derived from the onset of action of 20 nM STX at pH 7.3, l0°C from Fig. 10) was approximately 72 sec which may be compared to the mean value of 19 sec (pH 7.2, 16°C) obtained for STX block at the frog node of Ranvier (see Tables 2 & 3 of Wagner and Ulbricht, 1975). Given the differences in temperature and diffusion rates for the toxin at the node Ranvier and the squid giant axon (see Keynes et al., 1975), the only conclusion one can draw is that a slower association rate for STX was obtained under our experimental conditions.

Although there are similarities in the antagonism of toxin block of Nav channels by NH_4_^+^, N_2_H_5_^+^ and H^+^, the underlying mechanisms may differ. Perfusion with 300 mM NH_4_^+^ may lead to an alkalinization of the periaxonal space and thus lower the effective (protonated) toxin concentration. The two different experimental protocols to address this possibility were to examine the effect of internal perfusion on STX block: (1) with an elevated extracellular buffer concentration, and (2) in the presence of a lower external pH. The relief of STX block of Nav channels by internal NH_4_^+^ was observed, though reduced, during both experimental procedures, and is thus consistent with a mechanism of competition by NH_4_^+^ for a toxin binding site. In addition, comparison of NH_4_^+^ antagonism of STX and TTX (zwitterion with a pKa of 8.8) action indicates that the relief of toxin block is not due to a change in the degree of ionization of the toxin molecule. While these experiments lend support to a direct interaction between NH_4_^+^ and a toxin binding site, we cannot completely rule out the possibility that periaxonal pH changes play a role in this phenomenon. This question will perhaps remain unresolved until accurate measurements of pH changes in the periaxonal space during perfusion with NH_4_^+^ can be performed.

In summary, we have demonstrated an apparent trans channel interaction between the organic cations, NH_4_^+^ and N_2_H_5_^+^, in the internal compartment of the squid giant axon and STX or TTX molecules which bind to a site(s) on the external portion of the voltage-gated sodium channel. The apparent reduction in toxin potency observed under these conditions is similar to that seen by lowering internal pH. Our results support the notion that interaction of permeant cations with toxin binding sites may be more complicated than simple competition of ligands from the same cellular compartment.

## ACKNOWLEDGEMENTS

The authors thank Dr. Olaf Andersen for constructive suggestions during this project. This work was supported by National Science Foundation grant (BNS 8211580) to G.S. Oxford and a Wellcome Trust travel award to D.J. Adams.

The authors declare no competing financial interests.

Author contributions: G.S. Oxford, P Forscher, P.K. Wagoner and D.J. Adams designed and performed experiments, analyzed and interpreted data, and co-wrote the paper.

